# The benefits of estradiol on cognitive aging in rats are independent from its effects on cardiometabolic health

**DOI:** 10.1101/2024.12.10.627611

**Authors:** Christian Montanari, Emma L Dong, Shruti Srinivasan, Ana Paula De Oliveira Leite, Alyssa F Delarge, Matthieu J Maroteaux, Lucie D Desmoulins, Riva Menon, Alice B Walker, Sarah H Lindsey, Andrea Zsombok, Jill M Daniel

## Abstract

Research in preclinical models of menopause indicates that exogenously administered estrogens positively impact cognitive aging. However, clinical evidence indicates that the effects of estrogen therapy on cognition are inconsistent and may be modulated by pre-existing cardiometabolic conditions. The extent to which cardiometabolic health affects the cognitive outcomes of estrogen therapy remains unclear. This study aimed to determine whether variations in cardiometabolic health, both prior to and resulting from different estradiol treatment regimens, are related to the ability of estradiol to improve the cognitive aging trajectory in ovariectomized Long-Evans rats. Cognitive function and health status were assessed at 10 months of age after which rats were ovariectomized and administered vehicle or various estradiol treatments. Rats were assessed again at 18 (middle age) and 22 (old age) months. Cognition was evaluated using a spatial memory radial-maze task. Health status was determined through body composition (dual-energy X-ray absorptiometry), glucose tolerance testing, and blood pressure (tail-cuff plethysmography). Results demonstrated that both continuous ongoing estradiol treatment and a previous 40-day estradiol exposure (terminated long before testing) significantly improved the cognitive aging trajectory from middle to old age. However, only continuous estradiol treatment had positive impacts on health measures; previous estradiol treatment provided no benefits to aging cardiometabolic systems. In contrast, a delayed estradiol treatment (initiated months after ovariectomy) provided no benefits for cognition but provided health benefits. Results indicated that estradiol impacts on cognition in healthy aging rats are separate from and not secondary to its effects on cardiometabolic health.

## Introduction

Decades of research in preclinical models of menopause have consistently demonstrated that estrogens exert neuroprotective effects on the brain and improve memory in aging female rodents (1). Research has demonstrated that estrogens enhance synaptic plasticity (2), promote cholinergic neurotransmission (3), stimulate adult neurogenesis (4), and improve mitochondrial bioenergetics in neurons (5). These effects are particularly noticeable in brain regions critical for cognitive function, such as the hippocampus and cerebral cortex (1).

In women, the loss of ovarian estrogens at menopause is hypothesized to be a risk factor for Alzheimer’s disease (6–7). However, contrary to studies in animal models, clinical data have been inconsistent regarding the benefits of menopausal estrogen therapy on the brain and cognition, showing results that vary from beneficial to harmful (8–9). Early observational studies and small clinical trials suggested that hormone therapy might help prevent age-related cognitive decline (for review, see 10). Conversely, the large Women’s Health Initiative (WHI) Memory Study (11–14) conducted by the National Institutes of Health (NIH), reported that hormone therapy may increase the risk of dementia. The Kronos Early Estrogen Prevention Study (KEEPS, 15) and the KEEPS-Continuation Study (16) reported no harm or benefit from hormone therapy, as evaluated during the 4-year treatment period (15) and approximately 10 years after treatment termination (16).

Discrepancies in findings on the effects of estrogens on cognition between preclinical models and clinical studies, as well as among clinical studies themselves, might stem from differences in overall health status of subjects.

Indeed, preclinical research typically involves models of healthy aging, whereas clinical studies often include subjects with varying health conditions. Interestingly, preliminary evidence suggests that variations in clinical outcomes of hormone therapy might be partly explained by pre-existing cardiometabolic conditions, such as type 2 diabetes and hypertension, which may blunt the effects of estrogens on cognition (17–19).

The extent to which cardiometabolic health affects the cognitive outcomes of estrogen therapy remains unclear. In our laboratory, we have successfully established a surgical model of menopause in rodents and consistently demonstrated that midlife 17β-estradiol treatment in recently ovariectomized rats leads to long-lasting memory enhancements that persisted well beyond estradiol exposure (20–25).

The current study aimed to determine whether variations in cardiometabolic health, prior to and/or following different estradiol treatment regimens, influence the ability of estradiol to positively impact cognitive aging. We conducted a one-year longitudinal study assessing the cognitive aging trajectory and cardiometabolic health of rats from middle (10 months) to old (22 months) age. Our treatments aimed to model different populations of women, including those who start estrogen therapy at menopause and continue indefinitely, those who begin estrogen therapy at menopause but discontinue after a few years, those who start taking estrogens years after the onset of menopause, and those who never use menopausal estrogen therapy. We hypothesized that the effects of estrogens on cognition would be related to cardiometabolic health status. Understanding the nuanced relationship between cognitive and cardiometabolic health could lead to personalized treatment strategies that optimize cognitive health in aging females, ultimately reducing the risk of neurodegenerative diseases.

## Materials and Methods

### Subjects

Female Long-Evans rats (n=40) were obtained from Envigo at 70 days of age and allowed to age until the experiment began. The rats were pair-housed in a temperature-controlled vivarium on a 12-hour light/dark cycle (lights on at 7:00 a.m.), with unrestricted access to food (5V5R - PicoLab Select Rodent 50 IF/6F diet, PMI LabDiet, St. Louis, MO; it contains a targeted level of isoflavones for estrogen-sensitive protocols) and water, unless otherwise specified. All experimental procedures were performed during the light cycle. Animal care adhered to the National Institutes of Health’s Guide for the Care and Use of Laboratory Animals (26). All experimental procedures were approved by the Institutional Care and Use Committee of Tulane University. For experimental timeline, see Figure 1.

**Figure 1.**
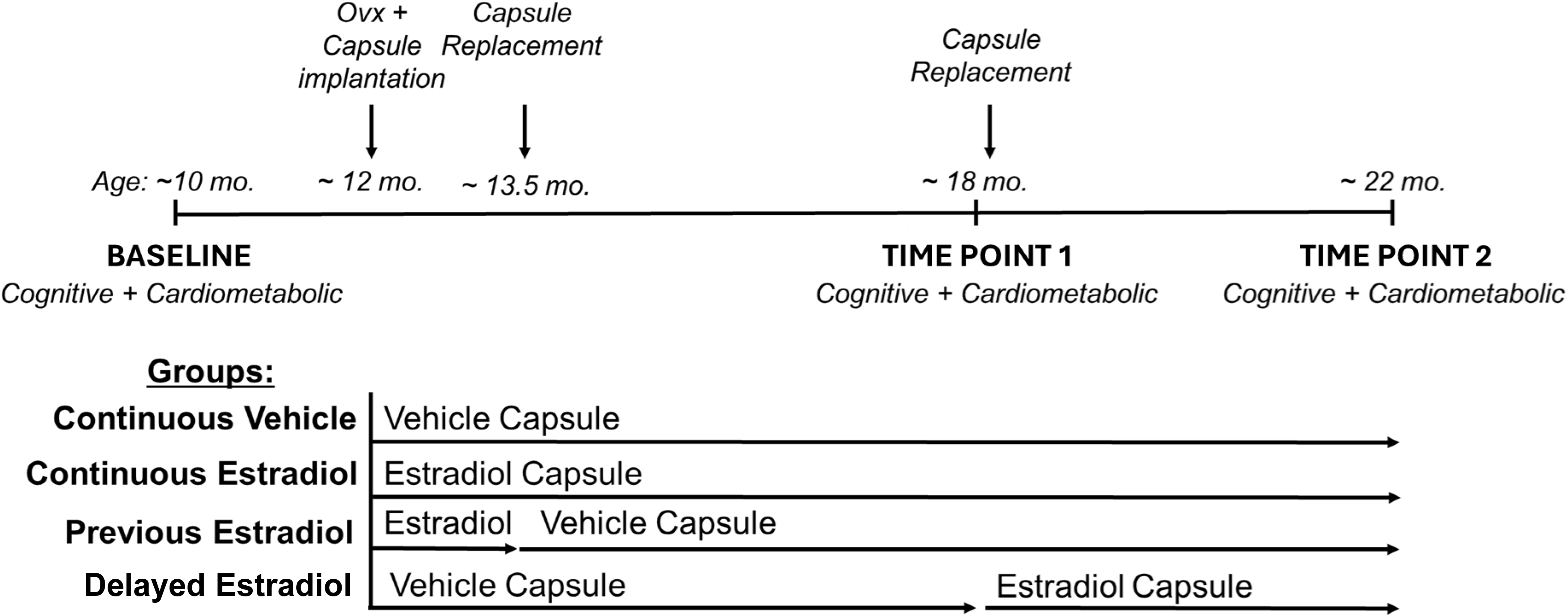
Experimental timeline. Cognitive function and cardiometabolic health were evaluated at Baseline (10 months of age). At 12 months of age, rats were ovariectomized and implanted with either estradiol or vehicle capsules. The capsules were replaced at 13.5 and 18 months of age, either maintaining the current vehicle or estradiol treatment or switching between them, resulting in the following treatment groups: Continuous Vehicle, Continuous Estradiol, Previous Estradiol, Delayed Estradiol. To assess the effects of the different estradiol treatments over time, cognitive function and cardiometabolic health were re-assessed at Time Point 1 (18 months of age, middle age) and at Time Point 2 (22 months, old age). Ovx, ovariectomy; mo., months.

### Radial-arm maze training and baseline cognitive assessment

When rats reached 9 months of age, training on the eight-arm radial maze spatial memory task began. During training, rats were food-restricted to maintain ∼90% of their free-feeding weight. Rats were trained for 24 days (one trial per day, 5 days per week). The maze (Coulbourn Instruments; 66 cm long, 9.5 cm wide, 11.5 cm high) had a metal grated floor and clear acrylic walls, with eight arms extending radially from a central hub (28 cm diameter). The maze was placed on a table ∼1 meter above the ground, centered in a room with multiple extra-maze cues. A single food reward (∼¼ of Froot Loops; Kellogg) was placed in an opaque dish (5.5 cm diameter, 1.25 cm tall) at the end of each arm, hidden from the maze center. For each trial, the rat was placed in the center of the maze, facing a pseudo-randomly chosen arm, and was allowed to explore the maze until all arms had been visited or 5 minutes elapsed. An arm entry was counted when all four paws crossed the arm midline, and re-entries (errors) were scored if the rat entered a previously visited arm. The arm entry sequence was scored in real time by an observer located in a fixed position in the room. Baseline cognitive function for each animal was determined at the end of training and when rats were 10 months of age by averaging the errors made during the first 8 arm choices over the final 4 days of radial-arm maze training.

### Baseline cardiometabolic health assessment

Following completion of radial-maze training, rats were returned to a free-feeding schedule. Approximately two weeks later and after body weights had stabilized at pre-training levels, baseline cardiometabolic assessment took place.

Cardiometabolic assessment included body composition analysis (dual-energy X-ray absorptiometry), glucose tolerance testing, and blood pressure measurement (tail-cuff plethysmography). The starting order of glucose tolerance and blood pressure tests were counterbalanced between subjects, with the body composition analysis being the final procedure.

*Dual energy X-ray absorptiometry (DEXA):* Body composition was determined using dual energy X-ray absorptiometry, calibrated with a reference phantom as described by the manufacturer (InAlyzer2, model S, Micro Photonics Inc., Allentown, PA, USA). Rats were anesthetized by gas inhalation (isoflurane 2%) during the whole procedure. After the animals were placed inside the main unit, DEXA scans were performed using the “rat” sample type with the “optimum” mode setting (scanning time <90s). Body mass, body fat, and body lean were collected as weight (in g).

*Glucose Tolerance Test (GTT):* After a 4-hour fasting period in clean cages with free access to water, rats were injected intraperitoneally with a solution of 20% glucose (2 g/kg body weight). Blood samples were collected from the tail vein before the glucose injection (T0, to assess basal glucose) and at 15-, 30-, 60-, and 120-minutes post-glucose injection to track changes in glucose levels over a 2-hour period and calculate its area under the curve (AUC). Blood glucose levels were measured using a glucometer (OneTouch Verio Flex).

*Tail-cuff plethysmography:* Systolic blood pressure (measured in millimeters of mercury, mmHg) was monitored using an automated tail-cuff volume-pressure recording system (Kent Scientific CODA® system). Animals were acclimated to the clear plastic tube restraints for 2 days and measurements were obtained over 3- 5 consecutive days to reduce the impact of restraint-induced stress. Ten to 15 consecutive measurements were taken for each animal while warming at 35°C under slight restraint. Measures were averaged over consecutive days excluding values that were ±2SD from the mean.

### Ovariectomy and hormone treatments

After completion of baseline cognitive and cardiometabolic assessments and when animals were approximately 12 months of age, experimental treatments began. All animals were ovariectomized. Rats were anesthetized with intraperitoneal injections of ketamine (100 mg/kg; Bristol Laboratories, Syracuse, NY) and xylazine (7 mg/kg; Miles Laboratories, Shawnee, KS) and administered buprenorphine (0.375 mg/kg subcutaneously; Reckitt Benckiser Health Care) prior to surgery. Ovariectomy surgery involved bilateral flank incisions through the skin and muscle wall and removal of ovaries.

At the time of ovariectomy, rats were implanted with a subcutaneous 5-mm Silastic capsule (0.058-inch inner diameter and 0.077-inch outer diameter; Dow Corning, Midland, MI) on the dorsal aspect of the neck. Capsules contained either cholesterol vehicle or 25% 17β-estradiol (Sigma-Aldrich, St. Louis, MO) diluted in vehicle. Forty days (at 13.5 months of age) and 6 months (at 18 months of age) after ovariectomy and initial capsule implantation, rats were anesthetized by gas inhalation (isoflurane 2%) for capsule replacement. Capsule replacement involved either maintaining the current vehicle or estradiol treatment, or switching between them, resulting in the following randomly assigned estradiol treatment regimens (see Figure 1): 1) Continuous Vehicle (n=10; received vehicle for 10 months and modeling women who never use menopausal estrogen therapy); 2) Continuous Estradiol (n=10; received estradiol for 10 months and modeling women who take and remain on estrogen therapy); 3) Previous Estradiol (n=10; received estradiol for 40 days followed by vehicle for 8.5 months to model women who take menopausal estrogen therapy only for a few years in midlife); 4) Delayed Estradiol (n=10; received vehicle for 6 months followed by estradiol for 4 months to model women who begin taking estrogens years after menopause). We have previously demonstrated that capsules remain active for up to five months (20) and maintain blood serum estradiol levels at approximately 37 pg/ml in middle-aged female Long-Evans rats (27), which is within the physiological range.

### Cognitive Aging Trajectory

After ovariectomies, rats underwent a weekly radial-arm maze testing trial, which was identical to the baseline trials, for the remainder of the experiment. During this time, they remained on free-feeding and underwent a 24-hour food deprivation period prior to each testing day. To determine the impact of the various hormone treatments on the cognitive aging trajectories of the animals, baseline performance that was determined before ovariectomy and hormone treatments was compared to performance at two later timepoints by averaging results (errors of first eight arm choices) from four testing trials at each time point. Time Point 1 assessment occurred when animals were approximately 18 months of age, which was 4.5 months after termination of estradiol treatment in the Previous Estradiol group and before estradiol treatment began in the Delayed Estradiol group. Time Point 2 testing occurred when animals were approximately 22 months of age, which was 4 months following initiation of estradiol treatment in the Delayed Estradiol group. Using the mean centering method, the distribution of errors averaged at Baseline was centered around zero by subtracting the mean of the data from each individual data point. The errors averaged at Time Point 1 and Time Point 2 were then used to assess the cognitive aging trajectory of rats in each group, calculated as the percentage change in the number of errors compared to baseline performance at 10 months (set at 0%).

### Cardiometabolic health assessment over time

Cardiometabolic evaluations (including body composition analysis, glucose tolerance testing, and blood pressure) were conducted at Time Point 1 (18 months) and Time Point 2 (22 months) approximately two weeks after completing spatial memory testing for the corresponding time point.

### Hormone treatment verification

Vaginal smears were collected from each rat for 4 consecutive days before each of the two capsule replacements (at approximately 13.5 and 18 months of age, respectively) to confirm ovariectomy and hormone treatments. Smears from ovariectomized, cholesterol-treated rats were characterized by a predominance of leukocytes, while smears from ovariectomized, estradiol-treated rats were characterized by a predominance of cornified and nucleated epithelial cells, indicating that hormone treatments were effective. At the time of euthanasia, a 1- cm sample of the right uterine horn was collected from each rat and weighed to verify hormone treatment at the time of death.

### Euthanasia and tissue collection

Rats were euthanized by decapitation under anesthesia induced by ketamine (100 mg/kg) and xylazine (7 mg/kg). Various organs and tissues (uterus, adrenal gland, kidney, spleen, liver, heart, inguinal fat, and visceral fat) were collected and weighed.

### Statistical Analysis

All variables are expressed as mean ± SEM. Statistical analyses and graphs were generated using GraphPad Prism 10.2.3 software. Significant main or interaction effects (p ≤ 0.05) were followed by Tukey post hoc tests.

Two rats were excluded for failing to perform the radial arm-maze task. Ten rats were euthanized at various times due to age-related health issues. Additional exclusions from GTT were due to procedural issues. Final sample sizes are provided in the figure legends.

Cognitive aging trajectory from 10 to 22 months of age, body composition parameters (body mass, body fat, body lean), basal glucose, glucose AUC and systolic blood pressure were assessed using a mixed-effects model analysis with treatment (Continuous Vehicle, Continuous Estradiol, Previous Estradiol, Delayed Estradiol) as the between-subjects factor and time (Baseline, Time Point 1, Time Point 2) as the within-subjects factor.

At each testing (Baseline, Time Point 1, Time Point 2), glucose levels following glucose injection were analyzed using two-way ANOVA, with treatment (Continuous Vehicle, Continuous Estradiol, Previous Estradiol, Delayed Estradiol) as the between-subjects factor and minutes post-glucose injection (0, 15, 30, 60, 120 minutes) as the within-subjects factor.

At both Time Point 1 and Time Point 2, Pearson’s correlation coefficient (two-tailed) was performed to investigate the relationship between cognitive aging trajectory (measured as the percentage increase from baseline in the number of errors in the radial arm maze) and cardiometabolic health measures. Correlation analyses were conducted on the total sample and independently within each treatment group, regardless of overall significance.

The weights of organs collected at the time of sacrifice were analyzed using one-way ANOVA.

## Results

### Cognitive aging trajectory

As illustrated in Figure 2, analysis of the percent change in number of errors during radial-arm maze testing at Time Point 1 and Time Point 2 compared to baseline performance (mean ±SEM: 0.91±0.073 errors), revealed significant main effects of treatment (F(3, 32) = 4.263, p = 0.0122) and time (F(1.810, 54.31) = 6.951, p = 0.0028) and no significant interaction of treatment × time (F(6, 60) = 2.027, p = 0.0758). Post-hoc comparisons of the significant main effect of treatment indicated that the cognitive aging trajectory of both the Continuous Estradiol and Previous Estradiol groups differed from both Continuous Vehicle and Delayed Estradiol groups (p ≤ 0.05 for all comparisons). Notably, the Continuous Estradiol and Previous Estradiol groups showed little to no aging-related decrement in performance as indicated by no change over time in the percentage of errors from baseline. In contrast, the Continuous Vehicle and Delayed Estradiol groups exhibited an increase of up to 125% in errors across time. Post-hoc comparisons of the significant main effect of time indicated that performance at Baseline across groups differed from performance at both Time Point 1 and Time Point 2 (p ≤ 0.05 for both comparisons).

**Figure 2.**
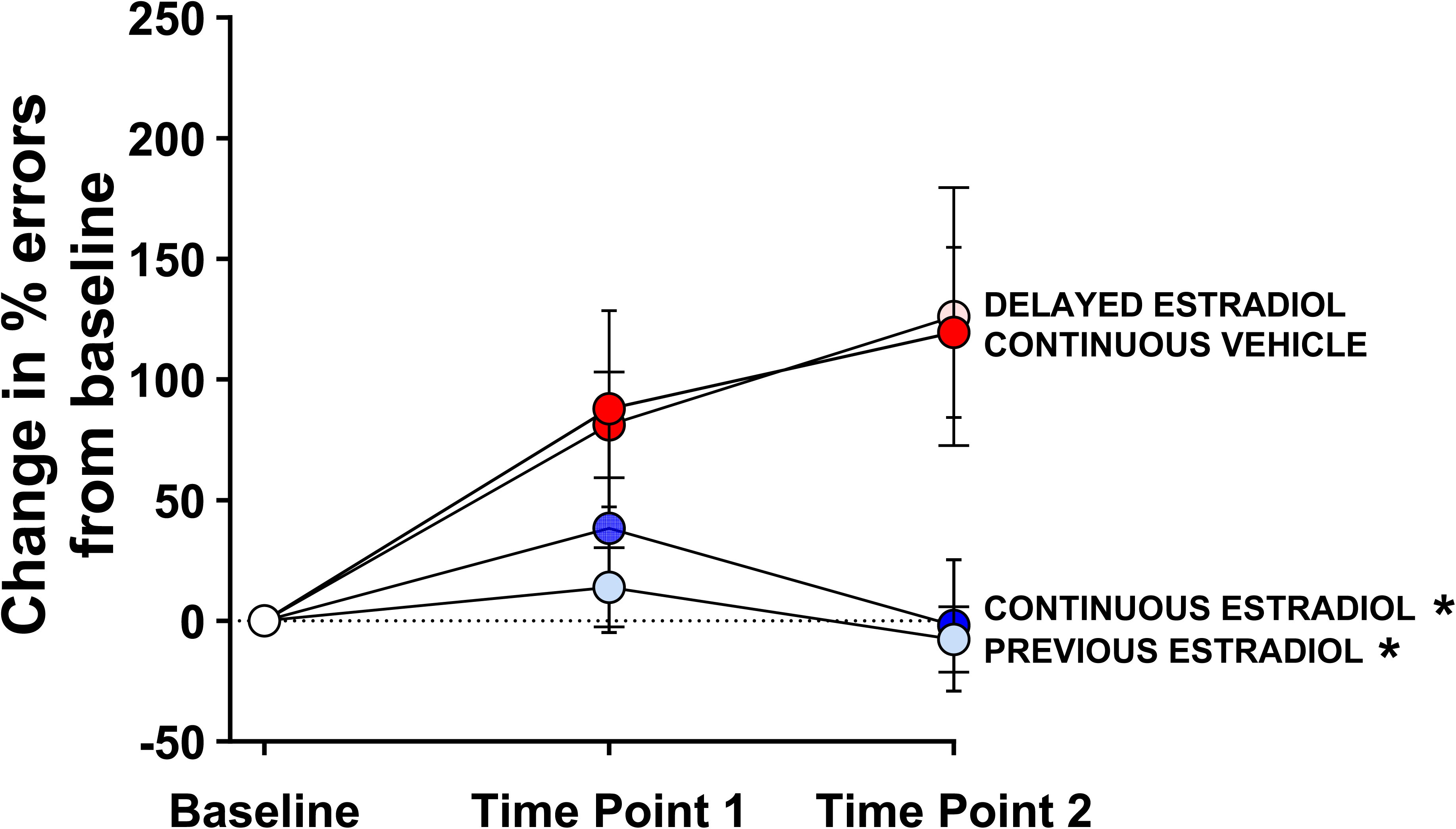
Cognitive aging trajectory. Mean (±SEM) percentage change in the number of errors during radial-arm maze testing at Time Point 1 (18 months of age) and Time Point 2 (22 months of age), compared to Baseline performance at 10 months, for the Continuous Estradiol, Continuous Vehicle, Previous Estradiol, and Delayed Estradiol groups. At Baseline, all groups are represented with white circles, as treatments had not yet begun. Following treatment initiation, groups were color-coded based on their respective estradiol treatment regimens: the Continuous Estradiol group is shown in blue circles at both Time Point 1 and Time Point 2 (indicating ongoing estradiol treatment at the time of testing); the Continuous Vehicle group is shown in red circles at both Time Point 1 and Time Point 2 (indicating ongoing vehicle treatment at the time of testing); the Previous Estradiol group is shown in light blue circles at both Time Point 1 and Time Point 2 (indicating estradiol treatment had been terminated months prior to testing); the Delayed Estradiol group is shown in red circles at Time Point 1 (indicating ongoing vehicle treatment at the time of testing), and in pink circles at Time Point 2 (indicating an ongoing delayed estradiol treatment initiated after testing at Time Point 1). Sample size at Baseline: Continuous Vehicle, n=8; Delayed Estradiol, n=8; Continuous Estradiol, n=10; Previous Estradiol, n=10; Sample Size at Time Point 1: Continuous Vehicle, n=8; Delayed Estradiol, n=8; Continuous Estradiol, n=10; Previous Estradiol, n=10; Sample size Time Point 2: Continuous Vehicle, n=7; Delayed Estradiol, n=8; Continuous Estradiol, n=7; Previous Estradiol, n=10. *p ≤ 0.05 compared to Continuous Vehicle and Delayed Estradiol.

### Body composition analysis

#### Body mass

As illustrated in Figure 3A, analysis of body mass revealed a significant main effect of treatment (F(3, 32) = 4.853, p = 0.0068) and time (F(2,1.171, 32.7856) = 161.2, p < 0.0001) as well as a significant interaction of treatment x time (F(6, 56) = 8.478, p < 0.0001). Post-hoc comparisons revealed no group differences at Baseline, before treatments began. However, at both Time Point 1 and Time Point 2, the Continuous Estradiol group exhibited significantly lower body mass compared to all other groups (p ≤ 0.05 for all comparisons). When evaluating changes in body mass over time, the Continuous Vehicle and Previous Estradiol groups showed higher body mass at both Time Point 1 and Time Point 2 compared to Baseline (p ≤ 0.0001 for all comparisons), with an additional increase from Time Point 1 to Time Point 2 (p < 0.05 for both groups). Results reveal a progressive weight gain in absence of estradiol (Continuous Vehicle) or when estradiol treatment was interrupted months before testing (Previous Estradiol). In the Delayed Estradiol group, body mass was significantly higher at both Time Point 1 and Time Point 2 compared to Baseline (p ≤ 0.0001 for both comparisons), but no difference was observed between Time Point 1 and Time Point 2 (p = 0.4383), indicating stabilization of body mass as estradiol treatment started after testing at Time Point 1. In contrast, the Continuous Estradiol group showed an increased body mass compared to Baseline only at Time Point 1 (p ≤ 0.01), suggesting that continuous ongoing estradiol treatment blunted naturally occurring weight gain following the loss of ovarian function.

**Figure 3.**
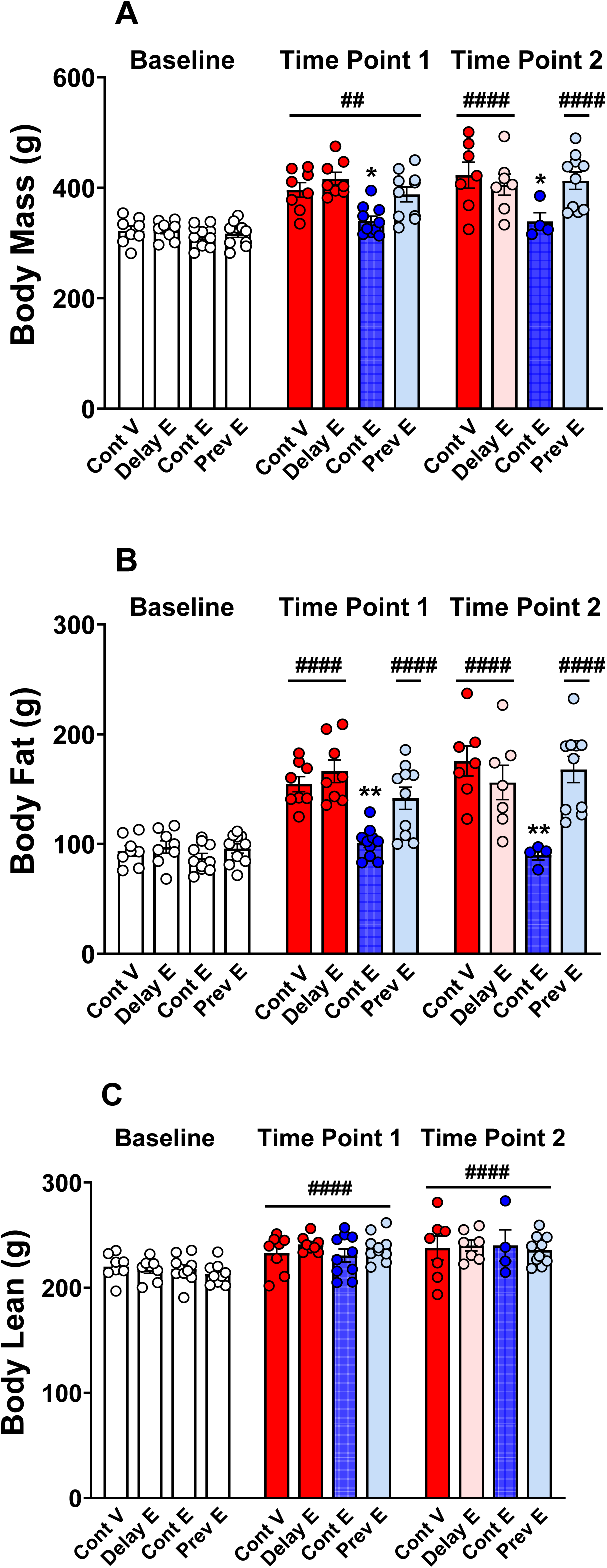
Body composition. Mean (±SEM) body mass (A), body fat (B), and body lean mass (C) measured at Baseline (10 months), Time Point 1 (18 months), and Time Point 2 (22 months) in the Continuous Vehicle (Cont V), Delayed Estradiol (Delay E), Continuous Estradiol (Cont E), and Previous Estradiol (Prev E) groups. At Baseline, all groups are represented with white bars, as treatments had not yet begun. At Time Point 1, the Delay E group is depicted by a red bar indicating ongoing vehicle treatment at the time of testing. Sample size at Baseline: Cont V, n=8; Delay E, n=8; Cont E, n=10; Prev E, n=10; Sample size at Time Point 1: Cont V, n=8; Delay E, n=8; Cont E, n=10; Prev E, n=10; Sample size at Time Point 2: Cont V, n=7; Delay E, n=7; Cont E, n=4; Prev E, n=10. *p ≤ 0.05, and ** p ≤ 0.01 compared to all the other groups; ## p ≤ 0.01, and #### p ≤ 0.0001 compared to the corresponding baseline within each group.

#### Body fat mass

As illustrated in Figure 3B, analysis of body fat mass revealed a significant main effect of treatment (F(3, 32) = 9.924, p < 0.0001), time (F(2, 56) = 125.1, p < 0.0001), and a significant time × treatment interaction (F(6, 56) = 12.32, p < 0.0001). Consistent with findings in body mass, no differences were identified among groups at Baseline (and before treatments began), whereas at both Time Point 1 and Time Point 2 the Continuous Estradiol group had lower body fat compared to the other treatment groups (p ≤ 0.01 for all comparisons).

Additionally, the Continuous Estradiol group did not show changes in body fat over time (Baseline vs. Time Point 1, p = 0.1106; Baseline vs. Time Point 2, p = 0.9789; Time Point 1 vs. Time Point 2, p = 0.4406). In the Continuous Vehicle and Previous Estradiol groups, body fat levels were elevated at both Time Point 1 and Time Point 2 compared to Baseline (p ≤ 0.0001 for all comparisons), with a further increase noted from Time Point 1 to Time Point 2 (p ≤ 0.05). For the Delayed Estradiol group, body fat was significantly higher at both Time Point 1 and Time Point 2 compared to Baseline (p ≤ 0.0001 for all comparisons), with no difference between Time Point 1 and Time Point 2 (p = 0.2829).

#### Body lean mass

As illustrated in Figure 3C, analysis of body lean mass indicated a significant main effect of time (F(2, 56) = 62.80, p < 0.0001), but no significant effect of treatment (F(3, 32) = 0.1343, p = 0.9389) or treatment × time interaction (F(6, 56) = 1.679, p = 0.1432). Post-hoc comparisons indicated that body lean was significantly higher at both Time Point 1 and Time Point 2 compared to Baseline (p < 0.0001 for both comparisons).

### Basal glucose levels

As illustrated in Figure 4, analysis of basal glucose levels prior to glucose injection revealed a significant main effect of treatment (F(3, 32) = 7.441, p = 0.0006) and time (F(1.672, 44.30) = 26.35, p < 0.0001) as well as a significant time × treatment interaction (F(6, 53) = 5.376, p = 0.0002). No group differences were detected at Baseline. At Time Point 1, basal glucose in the Continuous Estradiol group was lower than the Previous Estradiol group (p = 0.0201). At Time Point 2, basal glucose in the Continuous Estradiol group was lower than both the Previous Estradiol (p = 0.0007) and Continuous Vehicle (p = 0.0012) groups, while basal glucose in the Delayed Estradiol group was lower than the Continuous Vehicle group (p = 0.0275), indicating that ongoing estradiol treatment, even if initiated long past loss of ovarian function, lowered basal glucose levels. All groups except the Continuous Vehicle group showed lower basal glucose levels at Time Point 2 compared to Baseline (p ≤ 0.05 for all comparisons). A significant decrease between Time Point 1 and Time Point 2 was detected in the Delayed Estradiol and Previous Estradiol groups (p ≤ 0.01 for both comparisons).

**Figure 4.**
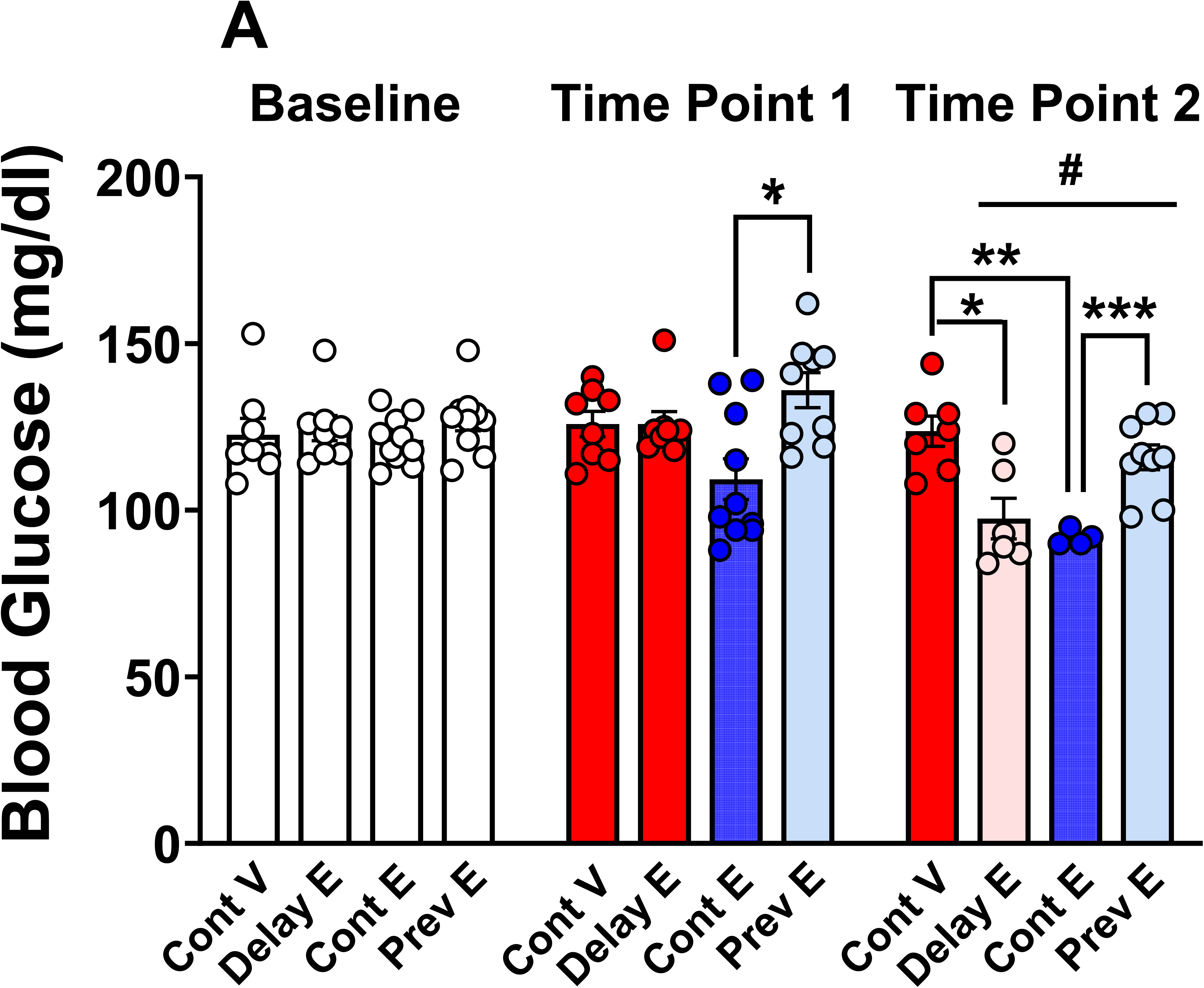
Basal glucose levels. Mean (±SEM) basal glucose levels at Baseline (10 months), Time Point 1 (18 months), and Time Point 2 (22 months) in the Continuous Vehicle (Cont V), Delayed Estradiol (Delay E), Continuous Estradiol (Cont E), and Previous Estradiol (Prev E) groups. At Baseline, all groups are represented with white bars, as treatments had not yet begun. At Time Point 1, the Delay E group is depicted by a red bar indicating ongoing vehicle treatment at the time of testing. Sample size at Baseline: Cont V, n=8; Delay E, n=8; Cont E, n=10; Prev E, n=10; Sample size at Time Point 1: Cont V, n=8; Delay E, n=8; Cont E, n=10; Prev E, n=9; Sample size at Time Point 2: Cont V, n=7; Delay E, n=6; Cont E, n=4; Prev E, n=9.*p ≤ 0.05, ** p ≤ 0.01, and *** p ≤ 0.001 indicate group differences; # p ≤ 0.05 compared to the corresponding baseline within each group.

### Glucose Tolerance Test

As illustrated in Figure 5, at each testing (Baseline, Time Point 1, Time Point 2), there was a significant main effect of minutes post-glucose injection (Baseline: F(3.175, 101.6) = 199.9, p < 0.0001; Time Point 1: F(2.315, 57.88) = 134.9, p < 0.0001; Time Point 2: F(2.162, 41.08) = 75.08, p < 0.0001). Post-hoc comparisons revealed that at each test, nearly all pairwise comparisons were significant (p ≤ 0.05 for all comparisons). Glucose levels at 15- and 30-minute post-glucose injection were significantly higher compared to basal pre-injection levels (0- minutes post-glucose injection; p ≤ 0.01 for all comparisons), while comparisons between 30- and 120-minute post-glucose injection indicated a decrease of glucose levels (p ≤ 0.05 for all comparisons). This suggests a peak in glucose levels 30 minutes after the glucose injection, followed by a gradual decline.

**Figure 5.**
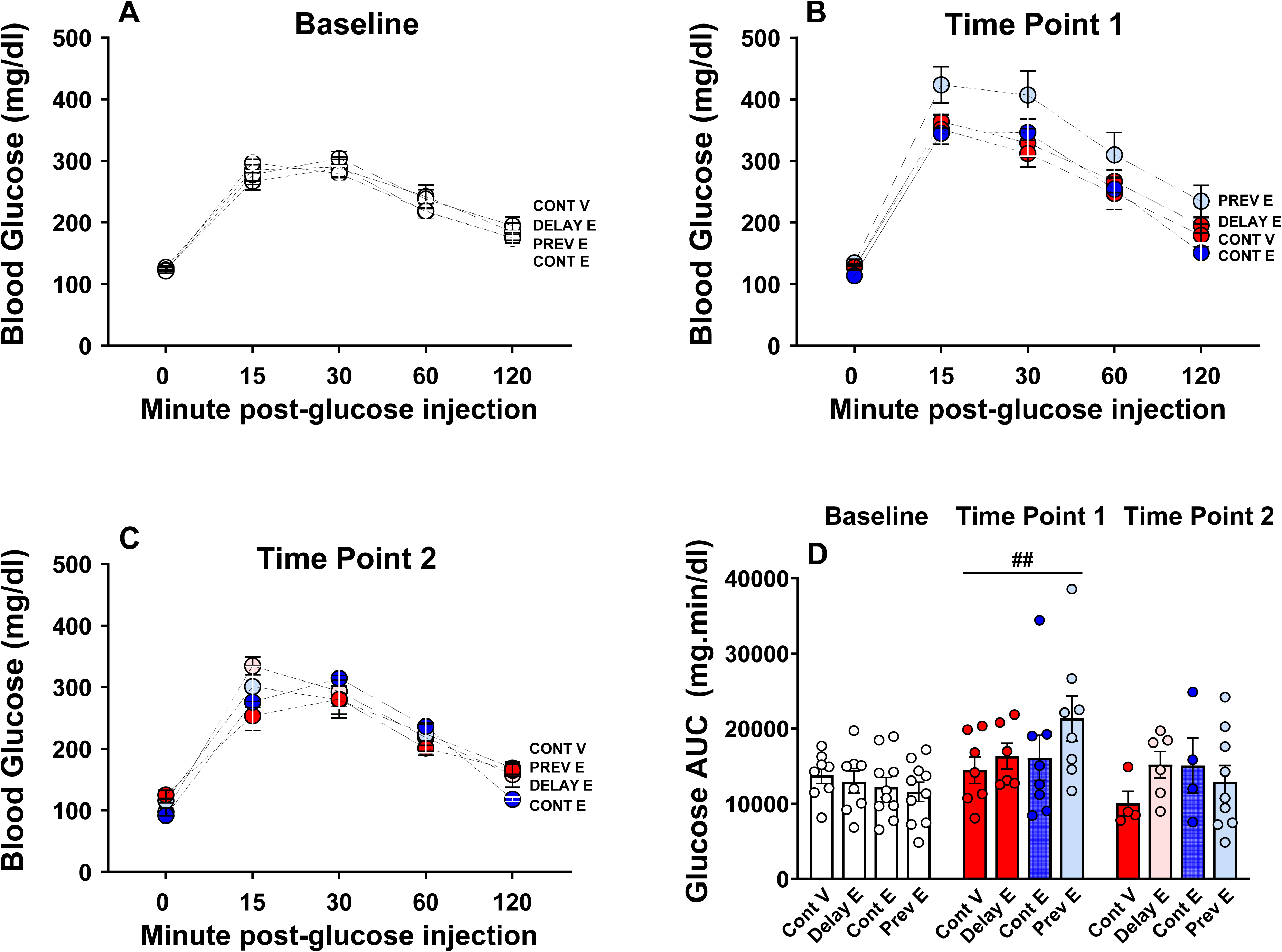
Glucose tolerance test. Mean (±SEM) glucose (A–C) levels over the 2- hour period following glucose injection measured at Baseline (A), Time Point 1 (B), and Time Point 2 (C) in the Continuous Vehicle (Cont V), Delayed Estradiol (Delay E), Continuous Estradiol (Cont E), and Previous Estradiol (Prev E) groups. The corresponding mean (±SEM) area under the curve (AUC) values are shown in panel D. At Baseline, all groups are depicted in white, as treatments had not yet begun. At Time Point 1, the Delay E group is depicted in red indicating ongoing vehicle treatment at the time of testing. Sample size at Baseline: Cont V, n=8; Delay E, n=8; Cont E, n=10; Prev E, n=10; Sample size at Time Point 1: Cont V, n=7; Delay E, n=6; Cont E, n=8; Prev E, n=8; Sample size at Time Point 2: Cont V, n=4; Delay E, n=6; Cont E, n=4; Prev E, n=9. ## p ≤ 0.01, compared to the corresponding baseline within each group.

No significant effect of treatment was found: Baseline: F(3, 32) = 0.1964, p = 0.8981; Time Point 1: F(3, 25) = 1.986, p = 0.1419; Time Point 2: F(3, 19) = 0.1531, p = 0.9264). There was no interaction of minutes post-glucose injection x treatment: Baseline: F(12, 128) = 1.122, p = 0.3478; Time Point 1: F(12, 100) = 0.9442, p = 0.5068; Time Point 2: F(12, 76) = 1.400, p = 0.1848).

Analysis of the glucose AUC revealed a significant main effect of time point (F(1.458, 55.40) = 5.535, p < 0.05). Post-hoc comparisons indicated a higher glucose AUC at Time Point 1 compared to Baseline (p = 0.0066). No significant main effect of treatment (F(3, 76) = 0.7635) or a significant time × treatment interaction (F(6, 76) = 1.392) was revealed.

### Systolic blood pressure

As illustrated in Figure 6, analysis of systolic blood pressure indicated no significant main effect of treatment (F (3, 32) = 1.672, p= 0.1927) or time (F (1.655, 44.68) = 3.113, p= 0.06331) but a significant treatment × time interaction (F (6, 54) = 3.085, p= 0.0114). Post-hoc comparisons showed a significant decrease in systolic blood pressure from Baseline to Time Point 2 in the Continuous Estradiol group (p = 0.0588). We also found a lower systolic blood pressure in the Delayed Estradiol group at Time Point 1 (prior to estradiol treatment initiation) compared to its Baseline (p = 0.0039).

**Figure 6.**
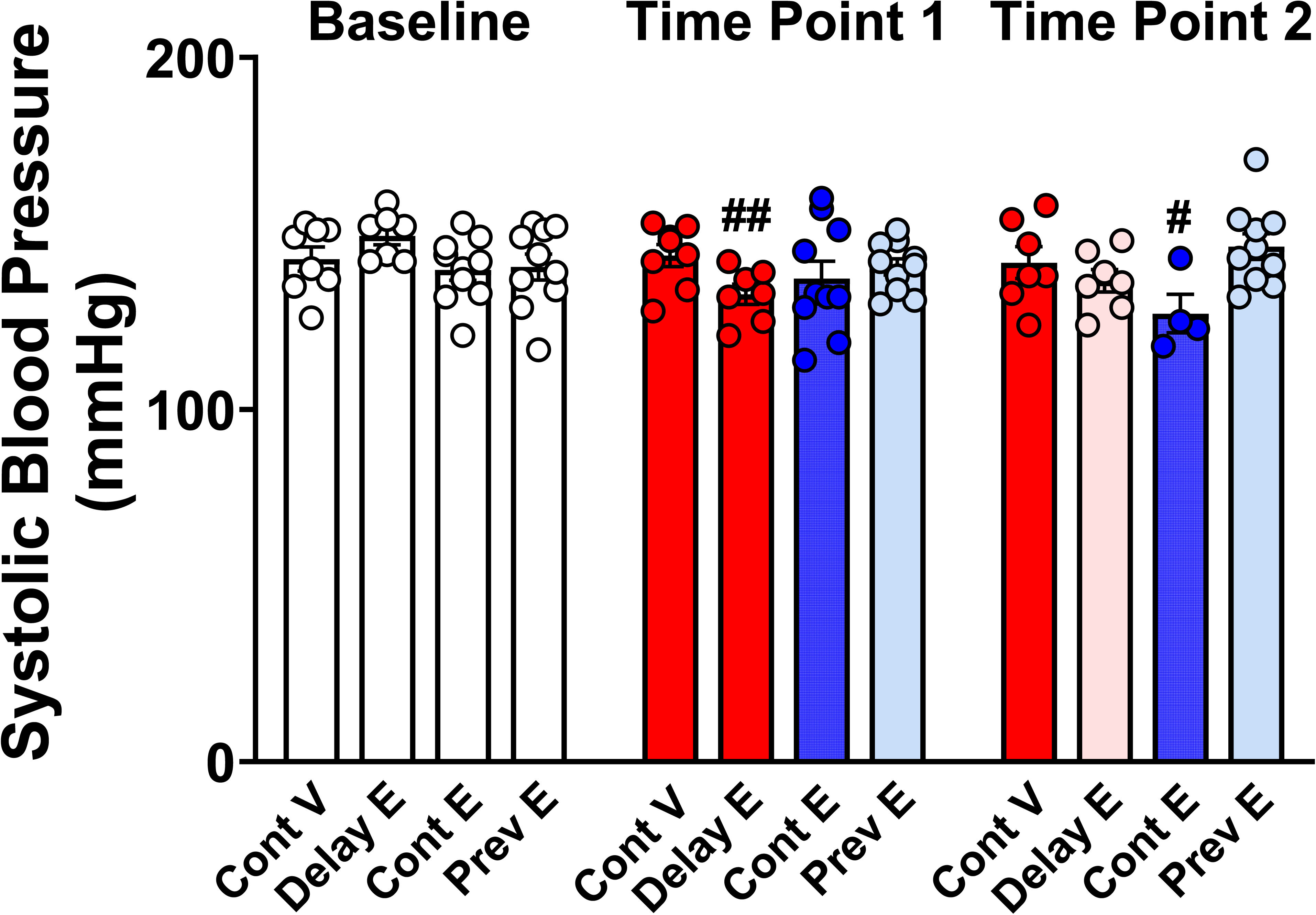
Systolic blood pressure. Mean (±SEM) systolic blood pressure measured in millimeters of mercury (mmHg) at Baseline (10 months), Time Point 1 (18 months), and Time Point 2 (22 months) in the Continuous Vehicle (Cont V), Delayed Estradiol (Delay E), Continuous Estradiol (Cont E), and Previous Estradiol (Prev E) groups. At Baseline, all groups are represented with white bars, as treatments had not yet begun. At Time Point 1, the Delay E group is depicted by a red bar indicating ongoing vehicle treatment at the time of testing. Sample size at Baseline: Cont V, n=8; Delay E, n=8; Cont E, n=10; Prev E, n=10; Sample size at Time Point 1: Cont V, n=8; Delay E, n=8; Cont E, n=10; Prev E, n=10; Sample size at Time Point 2: Cont V, n=7; Delay E, n=7; Cont E, n=4; Prev E, n=10. # p ≤ 0.05, and ## p ≤ 0.01 compared to the corresponding baseline within each group.

### Organ weights

As illustrated in Figure 7, there were significant effects of treatment for uterus (F(3, 24) = 4.326, p = 0.0142) and visceral fat (F(3, 24) = 5.199, p = 0.0066). Post-hoc comparisons indicated that the uterus weight of the Continuous Estradiol group was significantly higher than the Previous Estradiol (p = 0.0357) and the Continuous Vehicle (p = 0.0152) groups. For visceral fat, post-hoc comparisons showed that the Continuous Estradiol group had significantly lower visceral fat compared to the Continuous Vehicle group (p = 0.0044).

**Figure 7.**
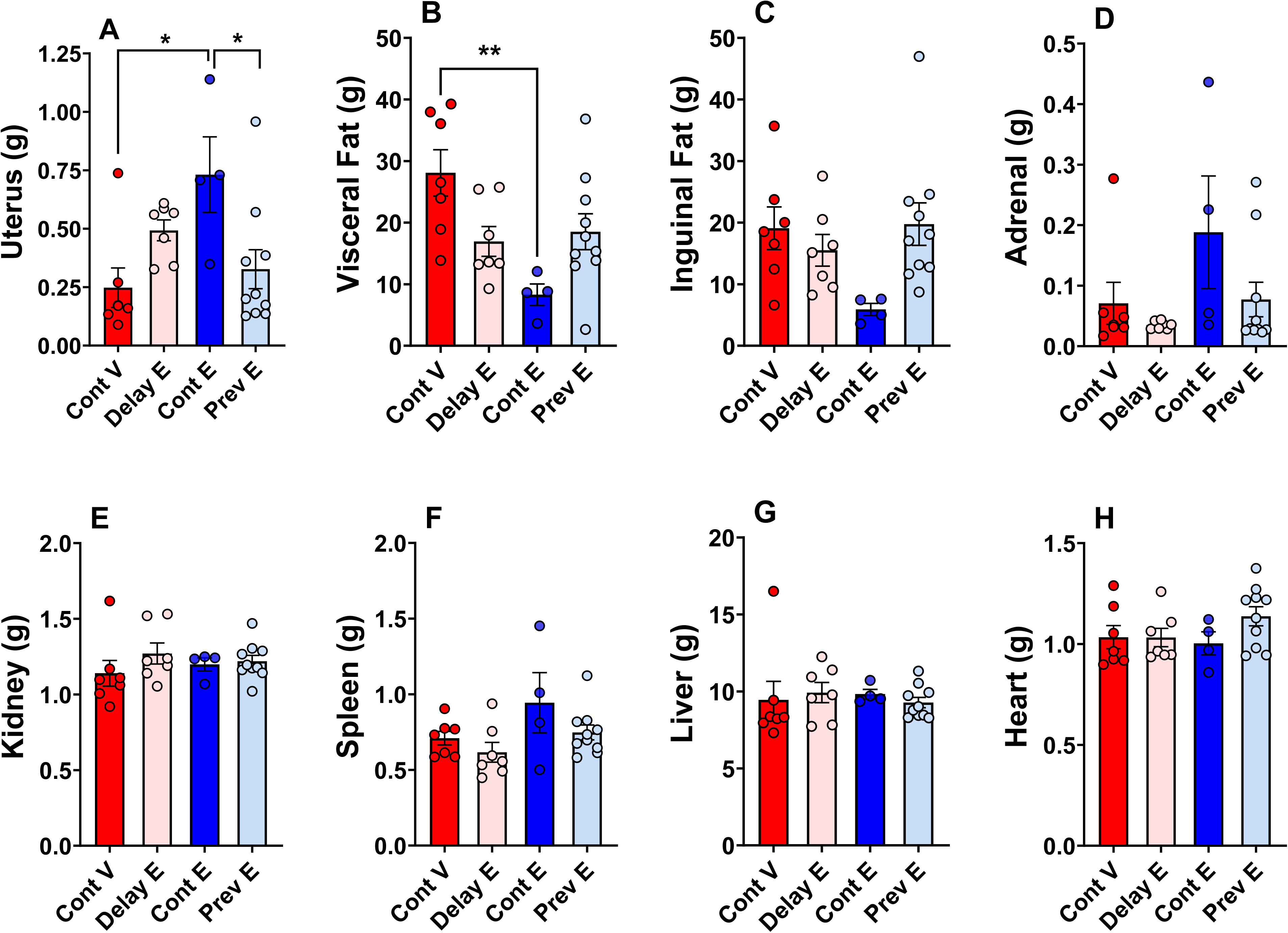
Organ weights. Mean (±SEM) weights of the uterus (A), visceral fat (B), inguinal fat (C), adrenal (D), kidney (E), spleen (F), liver (G), and heart (H) collected at 22 months of age for the Continuous Vehicle (Cont V), Delayed Estradiol (Delay E), Continuous Estradiol (Cont E), and Previous Estradiol (Prev E) groups. Sample size: Cont V, n=7; Delay E, n=7; Cont E, n=4; Prev E, n=10. *p ≤ 0.05 compared to Cont V and Prev E; ** p ≤ 0.01 compared to Cont V.

There were no significant treatment effects for other organs that were collected (inguinal fat: F(3, 24) = 2.728, p = 0.0746; adrenal: F(3, 24) = 2.159, p = 0.1191; kidney: F(3, 24) = 0.7455, p = 0.5355; spleen: F(3, 24) = 2.340, p = 0.0987; liver: F(3, 24) = 0.1889, p = 0.9029; heart: F(3, 24) = 1.411, p = 0.2640).

### Correlation Between Cognitive Aging Trajectory and Cardiometabolic Health Measures

As depicted in Table 1, correlation analyses were conducted across groups and independently within each treatment group, regardless of overall significance. Notably, only one significant correlation was identified: at Time Point 1, the cognitive aging trajectory (measured as the percentage increase in errors compared to Baseline) showed a significant negative association with systolic blood pressure (r =-0.3681, p = 0.0296). The negative correlation indicates that as systolic blood pressure increases, cognitive performance improves (i.e., a smaller percentage increase in errors). Subsequent analyses within each group revealed a significant correlation between cognitive aging trajectory and systolic blood pressure only in the Continuous Vehicle group (r =-0.7114, p = 0.0478). No significant correlations were observed in the Delayed Estradiol (r =-0.5954, p = 0.1584), Continuous Estradiol (r =-0.2756, p = 0.4409), or Previous Estradiol (r =-0.3244, p = 0.3605) groups.

**Table 1.**
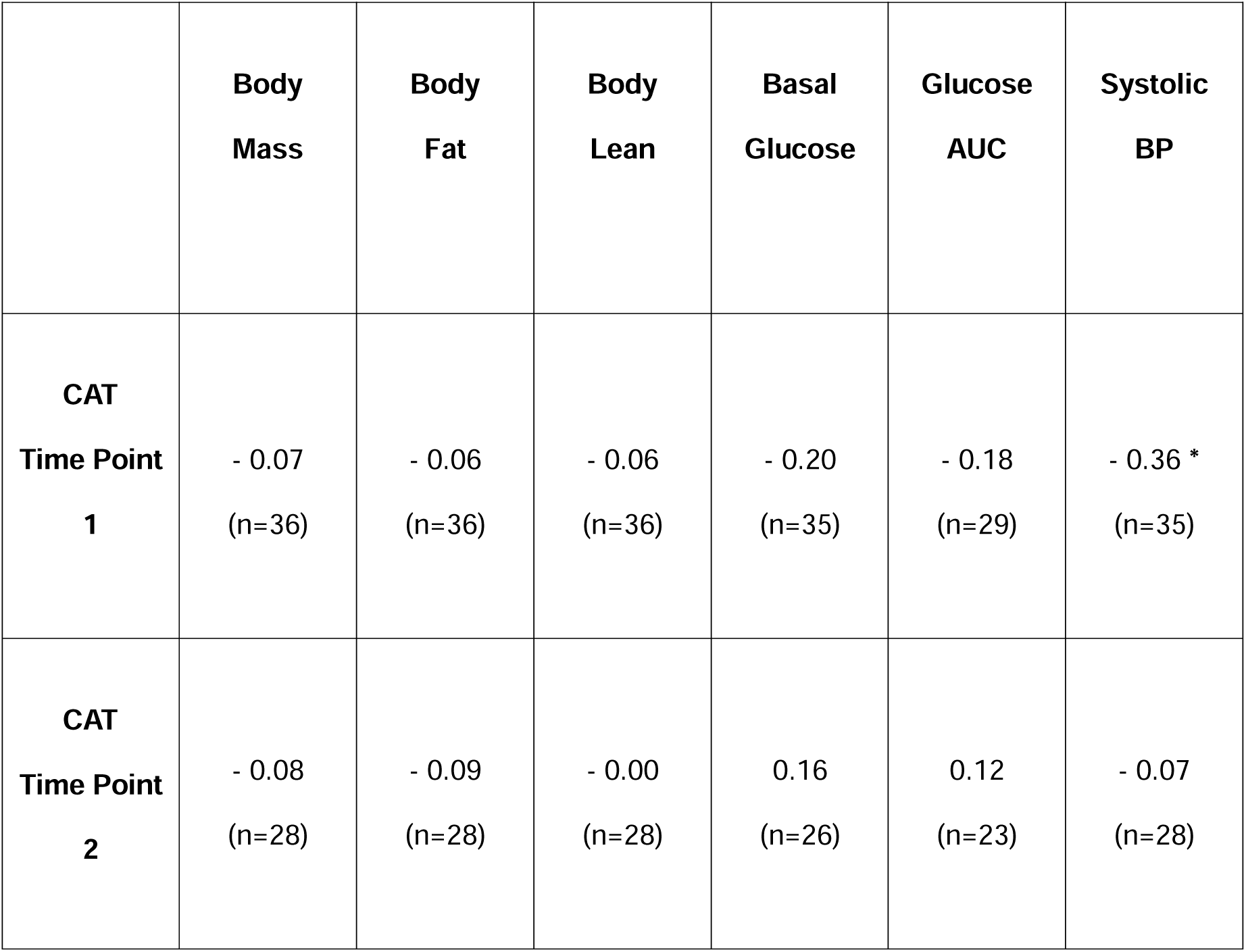
Correlation between cognitive aging trajectory and measures of cardiometabolic health. Pearson’s correlation coefficients (r) and corresponding sample sizes (n) for the relationship between cognitive aging trajectory and cardiometabolic health measures evaluated at both Time Point 1 and Time Point 2. CAT, Cognitive aging trajectory; AUC, Area under the curve; BP, Blood Pressure. * p ≤ 0.05.

No additional significant correlations were found between cognitive aging trajectory and any other cardiometabolic measures.

## Discussion

The current one-year longitudinal study followed aging ovariectomized female rats that were administered various regimens of estradiol treatments and assessed their cognitive aging trajectories and cardiometabolic health status from middle to old age (10–22 months of age). The goal of the work was to determine if there is a relationship between the ability of estrogens to impact cognitive aging and their ability to impact cardiometabolic aging in a rodent model of menopause. Results revealed that a regimen of continuous, ongoing estradiol treatment positively impacted both cognition (i.e. enhanced performance on spatial memory radial-arm maze task) and cardiometabolic health (i.e. reduction of body mass, body fat and visceral fat, basal glucose, and systolic blood pressure). In contrast, a previous history of 40 days of estradiol treatment that was terminated months prior to assessments had positive lasting effects on cognition, but no positive lasting impacts on measures of cardiometabolic health. Finally, a delayed estradiol treatment that was initiated six months after ovariectomy had no effect on cognition but had positive effects on some measures of cardiometabolic health (i.e. reduction of basal glucose and systolic blood pressure). Overall, our results indicate that estrogens actions on systems mediating cognitive function and those mediating cardiometabolic health are likely through independent mechanisms and that the impacts of estradiol on cognitive aging are not secondary to their effects on cardiometabolic systems.

### Effects of estrogens on cognition

The results of the present study indicate that, when initiated at the time of ovariectomy and subsequent ovarian function loss, both ongoing, continuous estradiol treatment (Continuous Estradiol) and a previous short-term estradiol treatment (Previous Estradiol), positively influenced the cognitive aging trajectory. This effect is observed in comparison to animals receiving no estradiol (Continuous Vehicle) or those that began estradiol treatment months after ovariectomy (Delayed Estradiol). Noteworthy, a previous 40-day estradiol treatment provided long-term cognitive benefits until the end of the experiment, corresponding to 10.5 months after estradiol treatment cessation. This effect was comparable to that observed in animals that were exposed to estradiol for the entire year of the experiment. On the other hand, exposing animals to estradiol 5 months after ovarian function loss did not benefit the cognitive aging trajectory, in spite of the duration of the delayed estradiol treatment of 4 months. These findings corroborate our previous results, showing that continuous estradiol treatment, as well as a prior estradiol exposure, significantly enhanced long-term memory compared to both a control group that never received estradiol and a group that started estradiol treatment months after ovariectomy (20–25, 27). However, unlike previous studies in which comparisons were made between groups, in the present study the assessment of the effects of estradiol on cognition was conducted over a year-long longitudinal study to track within each group the cognitive aging trajectory from middle to old age (10–22 months) and comparing cognitive performance to baseline levels prior to ovariectomy. Collectively, our results indicate that 1) estradiol treatment must be initiated within a critical window period after loss of ovarian function to confer cognitive benefits, and 2) a short 40- day estradiol regimen initiated at the time of ovariectomy offers long-term protection against cognitive decline, comparable to a continuous estradiol treatment.

Clinical data on the cognitive benefits of menopausal estrogen therapy have been inconsistent. Early observational and clinical studies (28-32; see 10 for a review) suggest that mid-life estrogen therapy following natural or surgical menopause provide long-term cognitive benefits. In contrast, the WHI Memory Study, a large randomized, double-blind, placebo-controlled clinical trial conducted by the NIH, found no cognitive benefit from hormone therapy. The study reported that hormone therapy regimens involving chronic conjugated equine estrogens (CEE) (13–14) or chronic CEE plus medroxyprogesterone (11–12) did not protect against age-related cognitive decline and may even increase the risk of dementia. However, unlike other studies, the WHI Memory Study administered hormone treatment to women aged 65 years and older (mean age of 73 years), which is, on average, more than two decades after ovarian hormone levels decline during menopause. Nonetheless, recent results from the KEEPS (15) and the KEEPS-Continuation investigations (16) found neither harm nor benefit from hormone therapy in recently postmenopausal women, keeping the debate still open. The KEEPS trial was a 4-year, randomized, double-blind, placebo-controlled trial designed to assess the effects of oral CEE or transdermal 17β-estradiol, both with progesterone, in healthy, recently postmenopausal women (mean age of 53 years, with an average of 1.4 years since menopause). The trial demonstrated no significant cognitive benefit or harm during the 4 years of menopausal hormone therapy (15). Moreover, to investigate the long-term effects of the previously administered treatments, the observational KEEPS-Continuation Study (16) evaluated cognitive outcomes approximately 10 years after completion of oral CEE plus progesterone or transdermal 17β-estradiol plus progesterone. Results revealed that women receiving either estrogen formulation exhibited similar cognitive performance to those assigned to placebo.

The critical period hypothesis has been proposed as a possible explanation for the discrepancies across studies, suggesting that the cognitive benefits of estrogens may only be apparent if treatment is initiated near the time of menopause (33–34); current clinical recommendations emphasize that hormone therapy provides the most benefit for patients when initiated within 10 years of menopause (North American Menopause Society, 2017, 35).

### Effects of estrogens on body composition

Our results indicate that ongoing, continuous estradiol exposure led to lower body mass and body fat compared to the other treatment groups. Moreover, unlike the other groups, body mass and body fat in the Continuous Estradiol group did not increase over time. In contrast to its lasting enhancement on cognitive aging, a previous history of 40-day estradiol exposure did not exert a lasting impact on body composition parameters. Indeed, the Previous Estradiol group showed similar body mass and body fat compared to the Continuous Vehicle and Delayed Estradiol groups, with both parameters increasing over time compared to baseline. Similarly, when estradiol treatment in the Delayed Estradiol group was initiated 5 months after ovariectomy, once significant body mass and fat augmentation had occurred, it was not able to suppress ovariectomy-induced body mass and fat increase, even with 4 months of treatment. Interestingly, the different estradiol treatment regimens did not affect body lean mass, which increased over time without significant group differences.

These findings are in line with our previous observations, in which ongoing estradiol treatment counteracted an increase in body mass following ovariectomy only if treatment was initiated at the time of ovarian function loss (3). In the present study we also evaluated changes in body fat and lean mass over a year-long period and weighed different organs at the time of sacrifice, highlighting that the ovariectomy-induced increase in body weight is primarily due to changes in the fat content. Moreover, continuous exposure to estradiol led to a reduction in visceral fat content by the end of the experiment.

Our findings align with human studies showing that changes in the hormonal milieu at menopause are associated with increased body mass and fat, which can be attenuated by estrogen administration (36). Moreover, studies have demonstrated a strong link between visceral fat and metabolic and cardiovascular disorders, with an increase in visceral adiposity identified as a risk factor for insulin resistance, type 2 diabetes, and cardiovascular disease mortality (37–38).

Our results indicate that estradiol regimens initiated at a healthy weight just after ovariectomy (Continuous Estradiol and Previous Estradiol groups) led to improved cognitive aging even under conditions of subsequent body weight and fat rise (Previous Estradiol). However, when estradiol was initiated months after ovariectomy and after body weight and fat increase occurred (Delayed Estradiol), no improvements in cognitive progression were observed. On one hand, these results support the critical period hypothesis, suggesting that the timing of estradiol initiation is crucial for its cognitive benefits. On the other hand, they also indicate that health status at the time of hormone treatment initiation may determine the efficacy of estradiol. These two possibilities are not mutually exclusive, and both the timing of hormone treatment initiation and health status at the time of initiation could synergistically contribute to the effects of estrogens on cognition.

### Effects of Estrogens on Glucose levels

In our study, ongoing estradiol treatment, initiated either at ovariectomy or months later, was associated with reduced basal blood glucose levels. Specifically, at 18 months of age (Time Point 1), the Continuous Estradiol group exhibited significantly lower basal glucose levels compared to the Previous Estradiol group, which had terminated treatment months prior. At 22 months (Time Point 2), the Continuous Estradiol group had lower basal glucose levels than both the Previous Estradiol and Continuous Vehicle groups, but not compared to the Delayed Estradiol group, which had significantly lower levels than the Continuous Vehicle group. These results indicate that ongoing estradiol treatment effectively reduces basal glucose, regardless of when treatment begins after ovarian function loss.

Congruent with these findings, previous preclinical studies support the critical role of estrogens in regulating glucose homeostasis and other metabolic parameters. Metabolic dysregulation has been observed in ovariectomized rats (39) and mice (40–41) on a standard diet. In these models, estradiol treatment has been shown to reduce both fasting and fed glucose levels, along with improvements in glucose tolerance (41). High-fat diet exposure is commonly used to induce obesity in rodent models, with female rats demonstrating greater protection against metabolic diseases; however, this sex-specific metabolic advantage diminished following ovariectomy (42).

Evidence from clinical trials indicates that estrogens treatment in non-diabetic postmenopausal women, started either in the early postmenopausal period (43–46) or within 10 years after menopause (47), is linked to reductions in fasting glucose and a lower risk of developing diabetes.

Although estradiol treatment resulted in significantly lower basal glucose levels, no group differences were observed in glucose levels during the GTT.

This divergence may reflect distinct regulatory mechanisms at play in basal versus glucose-stimulated conditions. Estradiol has been shown to enhance glucose uptake by upregulating and facilitating the translocation of glucose transporter 4 (GLUT4) to the cell surface (48), a process critical in maintaining basal glucose homeostasis. In contrast, other regulatory pathways may predominate during acute glucose challenges. For example, glucokinase, a central component of the glucostat system responsible for glucose homeostasis (49), plays a key role in liver-mediated glucose regulation by acting as a glucose sensor and initiating glycolysis in response to changes in blood glucose levels (50). This pathway integrates signals from various organs, such as the liver, pancreas, and brain, to maintain glucose homeostasis during feeding or glucose challenges (51).

### Effects of estrogens on cardiovascular systems

Our study showed that continuous estradiol treatment resulted in decreased systolic blood pressure at 22 months of age (Time Point 2), compared to its baseline (10 months). Studies conducted in rodent models indicate that experimentally induced ovarian function loss leads to decreased estradiol levels and an elevated risk of hypertension and cardiovascular diseases (for a review, see 52). Consistent with our findings in the Continuous Estradiol group, studies have shown that continuous estradiol treatment for 6 months (53) or 80 days (54) during a critical period post-ovariectomy, reduces systolic blood pressure in Long-Evans rats of similar age to those in our study.

As in rodents, menopause in humans leads to a decline in endogenous estrogens and an increased risk of cardiovascular diseases (52). Observational trials suggest that women receiving hormone therapy early in menopause may have a reduced risk of cardiovascular mortality later in life (55–57). Contrary to this evidence, the WHI Memory Study (58–59) found no cardiovascular benefits from estrogen therapy. Instead, it indicated that both chronic CEE and CEE plus medroxyprogesterone increased cardiovascular risk, irrespective of the timing of treatment initiation relative to menopause onset. However, a non-significant trend was observed, suggesting women who initiated hormone therapy closer to menopause might have a reduced risk compared to those more distant from menopause.

Our results also indicated a lower systolic blood pressure in the Delayed Estradiol group at 18 months (prior estradiol treatment initiation) compared to its baseline. This finding may be attributable to elevated systolic blood pressure at Baseline in this group, but not in the others, relative to the typical range for intact Long-Evans female rats of similar age (53–54).

### Relationship between the effects of estrogens on cognition and its effects on cardiometabolic systems

Numerous studies indicated that cognitive decline is more rapid in individuals with diabetes, who also face a higher risk of Alzheimer’s and related dementias than their non-diabetic peers (60). Interestingly, some studies suggested that certain antidiabetic medications, like metformin, reduce the risk of neurodegenerative diseases (61). In the women population, obese women with type 2 diabetes have twice the dementia risk of women at a normal weight (62). In two WHIMS follow-up studies, estrogen effects were evaluated within the context of type 2 diabetes.

One study found that estrogen alone - but not in combination with progestin - amplified diabetes-related risks for dementia and cognitive decline (18). Another study found that hormone therapy was linked to reduced brain volume in women with type 2 diabetes compared to those with type 2 diabetes who received a placebo, while brain volume in non-diabetic women was unaffected (63).

Numerous studies also demonstrated a direct association between cardiovascular complications and accelerated cognitive decline, contributing to Alzheimer’s disease and vascular dementia (64). The SPRINT MIND trial highlighted that while intensive blood pressure lowering (<120 mmHg) compared to standard treatment (<140 mmHg) does not prevent dementia, it does reduce the occurrence of mild cognitive impairment (65). Contrasting findings, such as those from the Hypertension in the Very Elderly Trial (HYVET), show that a year of antihypertensive treatment in adults over 80 years did not significantly lower dementia risk (66). However, age at intervention could explain these differences, as elevated systolic blood pressure after age 60 was counterintuitively found to be associated with improved cognitive performance (67). Regarding women in menopause, some findings indicated that initiating hormone therapy in the initial menopausal years reduces the risk of dementia and cardiovascular mortality years later (57).

Based on this evidence, our expectations for the present study were that cardiometabolic health would modulate the effects of estradiol on cognitive aging trajectory. Surprisingly, our results suggest that the benefits of estradiol on cognition are independent from its effects on cardiometabolic systems. Indeed, we found that both a regimen of continuous ongoing estradiol treatment and a prior short-term period of midlife estradiol treatment improved the cognitive aging trajectory from middle to old age. However, little to no relationship between the cognitive benefits of estradiol and its effects on cardiometabolic health was identified. For example, continuous estradiol treatment positively impacted both cognition and cardiometabolic health. In contrast, a previous history of 40 days of estradiol treatment terminated months prior testing had positive lasting effects on cognition, but did not confer any benefit to cardiometabolic health. Additionally, delayed estradiol treatment that was initiated months after ovariectomy showed no effect on cognition but had positive effects on some measures of cardiometabolic health. While our primary results indicate that the benefits of estradiol on cognition are not secondary to its effects on cardiometabolic systems, at 18 months of age we found a negative correlation between systolic blood pressure and cognitive performance, suggesting that as systolic blood pressure increases, cognitive performance improves. This counterintuitive finding aligns with recent human studies showing that elevated systolic blood pressure after age 60 is associated with reduced dementia risk (63).

## Conclusions

In conclusion, our study found no relationship between estradiol’s impact on cognitive aging and cardiometabolic health from middle to old age in healthy ovariectomized rats. These results highlight that the cognitive benefits of estradiol are not secondary to its cardiometabolic effects, suggesting that estradiol influences cognitive systems and cardiometabolic systems independently. This insight is significant because it challenges the view that cognitive improvements with estradiol may be a byproduct of cardiovascular and metabolic enhancements. Instead, our findings point to direct actions of estradiol on brain systems that support cognition, independent of metabolic and cardiovascular health, at least in the context of aging without pre-existing disease.

## Acknowledgments

The authors thank Dr. Janet Ruscher of the Department of Psychology at Tulane University for expert statistical advice.

## Data Availability

Datasets generated and analyzed during the current study are available from the corresponding author on reasonable request.

